# BioID analysis of the cyclin F interactome reveals that ALS-variant cyclin F alters the homeostasis of paraspeckle-associated proteins

**DOI:** 10.1101/2020.04.30.071365

**Authors:** Stephanie L. Rayner, Flora Cheng, Shu Yang, Natalie Grima, Yazi D. Ke, Carol G. Au, Marco Morsch, Alana De Luca, Jennilee M. Davidson, Mark P. Molloy, Bingyang Shi, Lars M. Ittner, Ian Blair, Roger S. Chung, Albert Lee

## Abstract

**Background:** Previously, we identified missense mutations in *CCNF* that are causative of familial and sporadic amyotrophic lateral sclerosis (ALS) and frontotemporal dementia (FTD). *CCNF* encodes for the protein cyclin F, a substrate recognition component of the E3-ubiquitin ligase, SCF^cyclin F^. We have previously shown that mutations in *CCNF* cause disruptions to overall protein homeostasis; causing a build-up of ubiquitylated proteins (*1*) as well as defects in autophagic machinery (*2*).

**Methods:** Here, we have used an unbiased proteomic screening workflow using BioID, as well as standard immunoprecipitations to identify novel interaction partners of cyclin F, identifying the interaction between cyclin F and a series of paraspeckle proteins. The homeostasis of these new cyclin F interaction partners, RBM14, NONO and SFPQ were monitored in primary neurons using immunoblotting. In addition, the homeostasis of RBM14 was compared between control and ALS/FTD patient tissue using standard IHC studies.

**Results:** Using BioID, we found over 100 putative interaction partners of cyclin F and demonstrated that cyclin F closely associates with a number of essential paraspeckle proteins, which are stress-responsive proteins that have recently been implicated in ALS pathogenesis. We further demonstrate that the turnover of these novel binding partners are defective when cyclin F carries an ALS/FTD-causing mutation. In addition the analysis of RBM14 levels in ALS patient post-mortem tissue revealed that RBM14 levels were significantly reduced in post-mortem ALS patient motor cortex and significantly reduced in the neurons of spinal cord tissue.

**Conclusion:** Overall, our data demonstrate that the dysregulation of paraspeckle components may be contributing factors to the molecular pathogenesis of ALS/FTD.

**Highlights:**

- Previously, we identified missense mutations in *CCNF* that are linked to Amyotrophic lateral sclerosis/Frontotemporal dementia (ALS/FTD) and have shown that a single mutation in cyclin F can cause defects to major protein degradation systems in dividing cells.
- Cyclin F has very few known interaction partners, many of which have roles in cell cycle progression. Accordingly, we used BioID and mass spectrometry to identify novel binding partners of cyclin F that may reveal insight into the role of cyclin F in neurodegeneration.
- Mass spectrometry and bioinformatic studies demonstrate that cyclin F interacts with several RNA binding proteins. This includes the essential paraspeckle proteins, RBM14. Notably, this interaction could be validated by standard immunoprecipitations and immunoblotting. Cyclin F could also be found to interact with a series of essential proteins which form the paraspeckle complex.
- We further evaluated the effect of cyclin F(S621G) on the homeostasis of these novel interaction partners in primary neurons in response to a known paraspeckle inducer, MG132. Notably, we demonstrate significant defects in the homeostasis of RBM14 and SFPQ, but not NONO, when cyclin F carries an S621G mutation.
- Unlike other paraspeckle proteins, RBM14 levels have not previously been reported in the post-mortem brain and spinal cord of ALS patient post-mortem tissue. Here, we note significant defects in the homeostasis of RBM14 in the post-mortem tissue of ALS patients.

## Background

Amyotrophic lateral sclerosis (ALS) is typically a late-onset neurodegenerative disease characterised by the selective degeneration of upper and lower motor neurons of the cerebral cortex, brainstem and spinal cord. It is the most common form of motor neurone disease (MND) with poor prognosis and limited treatment options. A proportion of ALS patients also develop clinical or subclinical frontotemporal dementia (FTD) and pathological and genetic overlap is now recognised, indicating that they represent a spectrum of disease (*3*). Approximately 5-10% of ALS patients carry an autosomal dominant genetic mutation. Familial mutations have been reported in over 30 genes including *SOD1* (*4, 5*), *VCP* (*6*), *TARDBP* (*7*), *FUS* (*8, 9*), *OPTN* (*10*), *SQSTM1* (*11*), *UBQLN2* (*12*), *MATR3* (*13*) and *TBK1* (*14, 15*). Identification of these genes has drawn attention to protein clearance pathways, proteins that accumulate within insoluble cytoplasmic inclusions and defects in RNA processing in disease pathogenesis. Recently, we identified several novel missense mutations in *CCNF* in patients with ALS/FTD (*1*). *CCNF* encodes for cyclin F, a 786 amino acid protein that forms part of the multi-protein Skp1-Cul1-F-Box (SCF^cyclin F^) E3 ligase that is known to regulate cell cycle progression through timely ubiquitylation of substrates to regulate their homeostasis through proteasomal degradation (*16*).

We have previously reported that a familial ALS/FTD mutation in cyclin F (denoted cyclin F^S621G^) alters the ubiquitylation activity of SCF^cyclin F^, leading to the accumulation of ubiquitylated proteins (*1*). In addition, we have also shown that the activity of cyclin F may be regulated by post-translational modifications and that the loss of a phosphorylation site causes aberrant ubiquitylation activity (*2*). Ultimately this leads to defects in bulk degradation processes and an upregulation in caspase-mediated cell death pathways (*17*).

Currently, there are few known interaction partners of cyclin F. These proteins are generally associated with cell-cycle function, including substrates such as ribonucleoside-diphosphate reductase subunit M2 (RRM2) (*18*), nucleolar and spindle-associated protein 1 (NuSAP) (*19*), centriolar coiled-coil protein of 110 kDa (CP110) (*20*), cell division control protein 6 homolog (CDC6) (*21*), stem-loop binding protein (SLBP) (*22*), exonuclease 1 (exo1) (*23*) and fizzy-related protein homolog (Fzr1) (*24*). In addition, known interaction partners of cyclin F include Skp1 (forming part of the ubiquitin ligase complex), b-myb (*25*) and CKII (*26*). Given that the interaction partners of cyclin F that have been reported to date are predominantly involved in cell-cycle regulation, it is not immediately obvious how cyclin F^S621G^ might trigger neurodegeneration in non-dividing neurons. Therefore, we hypothesised that there are other interaction partners of cyclin F that may help to understand the processes that may become defective in non-dividing cells.

BioID can be used to identify protein interaction partners and proteins in close proximity (∼10 nm radius) (*27*) to the protein of interest using an engineered biotin-ligase, BirA* (*28, 29*). An advantage of using BioID over standard immunoprecipitation (IP) methods, is the ability to identify transient, low abundance interaction partners as well as proteins that are not soluble in standard IP buffers (*30*). In recent years, BioID has been used to identify novel binding partners of a number of proteins including lamin A (*31*) and E-cadherin (*32*), ZO-1 (*33*), TDP-43 and fragmented TDP-43 (*34*). In addition, BioID has been utilized to identify substrates of β-TrCP 1 and 2 (*35*).

In this study, we have used BioID followed by mass spectrometry (MS) to characterise the interactome of cyclin F. In doing so, we identified more than 100 putative interaction partners of cyclin F, including a group of RNA binding proteins that are also essential paraspeckle proteins. Previously we have demonstrated that an ALS-causing mutation in cyclin F causes defects in major protein degradation systems, thus we evaluate the dysregulation of these proteins in primary neurons and in ALS patient tissue.

## METHODS

### Plasmids and Cloning

Expression constructs encoding wild type and S621G *CCNF* cDNA fused to an N-terminal mCherry fluorophore were used as described previously (*1*). Wild type and S621G *CCNF* cDNA fused to a C-terminal Flag-tag was also cloned into a pcDNA 3.1 vector. BirA* alone or BirA* in frame with cyclin F was cloned into pcDNA5/FRT/TO. Constructs encoding RBM14-HA were cloned into a pcDNA3.1 vector.

### Cell culture

Human Embryonic Kidney Cells (HEK293) and HEK293 Flp-In T-Rex cells were grown and maintained in Dulbecco’s modified Eagle medium (DMEM, Sigma-Aldrich) supplemented with 10% (v/v) of heat-inactivated fetal bovine serum (FBS, Sigma-Aldrich). Plated cells were grown and maintained in a humidified incubator held at a constant temperature of 37°C, with 5% CO_2_. All cell lines were tested for mycoplasma prior to experimental work using the MycoAlert Mycoplasma Detection Kit (Lonza).

In order to generate stably-transfected cell lines, HEK293 Flp-In T-Rex cells (Thermo) were double-transfected using constructs encoding Flp-recombinase (pOG44) as well as constructs encoding BirA*-cyclin F using Lipofectamine 2000 (Thermo) according to the manufacturer’s instructions. After 48 hours, cells were selected with 100 µg/mL Hygromycin (InvivoGen) and 15 µg/mL Blasticidin (InvivoGen). In order to ensure the cells were stably-transfected and that transgene expression could be induced, tetracycline (Sigma-Aldrich) was added to cell culture media at a final concentration of 0.1µg/mL for 18-24 hours. Tetracycline-dependent gene expression was monitored using standard immunoblotting procedures.

### Primary cell culture

Primary mouse cortical neurons were cultured as previously described (*36*). Briefly, brains were obtained from embryos on embryonic day 16.5. Cerebral hemispheres were sub-dissected, digested in trypsin at 37°C and homogenized using fire-polished glass pipettes into single cell suspension. Cells were seeded out at 5 million cells per 10 cm dish in medium containing 10% FBS/high glucose DMEM (Life Technologies). Medium was changed 2 hours post seeding and cells were subsequently maintained in Neurobasal medium supplemented with Glutamax and B27 supplement (Life Technologies).

### Proximity-labelling in live HEK293 Flp-In T-Rex cells

Stably transfected HEK293 Flp-In T-Rex cells were grown and maintained in DMEM supplemented with 10% FBS and 100 µg/mL Hygromycin (InvivoGen) and 15 µg/mL Blasticidin (InvivoGen). Once cells reached 70% confluency, expression of BirA* or BirA*-cyclin F (wild-type and *CCNF* variants) was induced by adding 0.1 µg/mL of tetracycline (Sigma-Aldrich) to cell culture media. In order to biotinylate proteins in proximity to the transgene, 50 µM of biotin (Sigma-Aldrich) was simultaneously added to the culture media. After 18-24 hours, cells were washed with PBS and harvested into ice-cold PBS. Harvested cells were washed twice with ice-cold PBS and centrifuged at 2000×g for 10 minutes at 4°C. Washed cell pellets were snap frozen at −80°C until further use.

### Total cell lysis

For total cell lysis, frozen cell pellets were first defrosted on ice and resuspended in ice-cold modified RIPA buffer (50 mM Tris-HCl, 150 mM NaCl, 1% NP-40, 1mM EDTA, 1mM EGTA, 0.1% SDS, 0.5% Sodium deoxycholate, pH 7.4) containing appropriate amounts of protease and phosphatase inhibitor cocktails (Roche). Cells were incubated in RIPA buffer for 15 minutes on ice with intermittent vortexing before probe sonication using a Sonic Ruptor 250 at 50% power and pulser settings set to 30%. Lysates were subject to a total of 10 pulses each before centrifugation at 14,000×g for 20 minutes at 4°C. The supernatant containing cellular proteins was aliquoted and stored at −80°C until further analysis.

### Biotin pull-downs

Cleared lysates containing biotinylated proteins in modified RIPA buffer were incubated with 30 µL of pre-washed streptavidin-coated magnetic beads (Thermo Fisher) for 3 hours at 4°C whilst rotating. In order to isolate biotinylated proteins from the complex mixture, a magnetic rack was used to isolate magnetic beads. Isolated magnetic beads were washed 5 times in modified RIPA buffer. Captured biotinylated proteins were eluted by resuspension in Laemmli sample buffer (BioRad), containing NuPAGE Sample Reducing Agent (Invitrogen) and were boiled at 95°C for 10 minutes. The eluents were prepared for 1D SDS-PAGE as described below.

### Immunoprecipitations

HEK293 cells were transfected with constructs encoding mCherry-cyclin F, Flag-cyclin F or RBM14-HA using Lipofectamine 2000 according to the manufacturer’s instructions. Transfected cells were harvested after 24 hours and cell pellets were resuspended in NP-40 lysis buffer (1 % (v/v) Nonidet P-40 in Tris-buffered saline (TBS), 2 mM EDTA, cOmplete protease inhibitor cocktail and phosSTOP (Roche)). The resuspended cells were vortexed, then probe sonicated (10 seconds, Setting 3, Branson Sonifier 450). The cell lysates were centrifuged at 14, 000×g for 30 minutes to remove cell debris. For immunoprecipitations, approximately 500 µg of cellular protein was incubated with 1 ug of flag antibody or 20 µL of RFP-Trap®_MA (Chromotek). The magnetic beads were collected using a magnet and washed three times in NP-40 lysis buffer. For western blot analysis, beads were resuspended in 1x Loading buffer (BioRad) containing 1x reducing reagent (NuPage) and boiled at 95°C for 10 minutes.

### SDS PAGE and Immunoblotting

Equal amounts of protein were separated on a 4-12% Bis-Tris SDS PAGE gel. Proteins were transferred onto a nitrocellulose membrane using a Bio-Rad Trans-blot Turbo semi-dry transfer cell (1.3 A, 25 V, 7 mins). The membranes were blocked in 3% skim milk powder in PBST for half an hour prior to incubation with primary antibody overnight at 4°C or 1 hour at RT. Primary antibodies used in this study were: rabbit polyclonal anti-cyclin F (1:300; cat# sc-952, Santa Cruz Biotechnology), mouse monoclonal anti-mCherry (1:300; cat# 632543, Clonetech), mouse monoclonal anti-β-actin (Abcam, dilution-1:12,000, catalogue #ab6276-101), mouse monoclonal anti-GAPDH (Proteintech, dilution-1:10,000), mouse monoclonal anti-α-tubulin (Sigma-Aldrich, dilution-1:1000, catalogue #T5168), rabbit polyclonal anti-RBM14 (Sigma-Aldrich, dilution-1:1000, catalogue #HPA006628), mouse monoclonal anti-PSPC1 (Santa Cruz, dilution-1:500, catalogue #sc-374367), mouse monoclonal anti-PSF (Santa Cruz, dilution-1:1000, catalogue #sc-101137), rabbit polyclonal anti-Matrin 3 (Proteintech, dilution-1:1000, catalogue #12202-2-AP).

After incubation with primary antibodies, the membranes were washed in PBS-T three times for 10 minutes before fluorescently labelled IRDye 800CW Goat Anti-Rabbit IgG Secondary Antibody (1:15,000; LI-COR) or fluorescently labelled IRDye^®^ 680RD Goat anti-Mouse IgG Secondary Antibody (1:15,00; LI-COR) secondary antibodies was added for 30 minutes at RT. Immunoblots were imaged using a Li-Cor Odyssey imaging system at the appropriate wavelength.

### In-gel trypsin digestion

Equal amounts of protein were loaded and separated on a 4-15% SDS-PAGE gel (BioRad). The resulting gel was briefly incubated in fixing solution (50% methanol, 10% acetic acid) and proteins were stained with Coomassie blue R250 until protein bands were visible. The gel was then left to destain overnight in Destain solution (25% methanol). After destaining, protein bands were excised from gels into 5 fractions. Gel fractions were then cut into smaller pieces (∼1 mm^2^) and further destained with 50% methanol/50 mM ammonium bicarbonate (pH 8). Gel pieces were then washed and dehydrated in 50% acetonitrile (ACN)/50 mM ammonium bicarbonate for 10 minutes, then incubated with 100% ACN until gel pieces were completely dehydrated. ACN was removed, and gel pieces were dried under vacuum centrifugation before being incubated with 10 mM dithiothreitol (DTT) in 50 mM ammonium bicarbonate (AmBic) for 40 minutes at 37°C.

Excess DTT was removed before gel pieces were incubated with 25 mM iodoacetamide (IAA) in 50 mM ammonium bicarbonate for 40 minutes at room temperature in the dark. Gel pieces were then washed twice with 50% ACN/50 mM ammonium bicarbonate for 10 minutes each time before the supernatant was removed and gel pieces were incubated in 100% (v/v) ACN to dehydrate gel pieces as described earlier. Excess ACN was removed and gel pieces were left to dry.

Gel pieces were incubated with trypsin (12.5 ng/µl; proteomics grade, Sigma-Aldrich) diluted in 50 mM ammonium bicarbonate and incubated overnight at 37°C. After incubation, the supernatant was transferred into fresh tubes and acidified with formic acid (FA). The gel pieces were incubated in 50% ACN, 2% FA. Supernatants containing tryptic peptides were pooled and lyophilised. For desalting, peptides were resuspended in 0.1% FA and desalted using pre-washed and equilibrated C18 OMIX tips (Agilent). Once desalted, samples were again lyophilised and stored at −80°C until MS analysis.

Prior to mass spectrometry, lyophilised peptides were resuspended in 0.1% FA and bath sonicated for 20 minutes. The resuspended peptides were then centrifuged at 14, 000×g for 15 minutes to remove any insoluble debris, and the clarified peptides were analysed by LC-MS/MS. The peptide fractions were separated on an Ultimate 3000 nanoLC (Thermo Fisher Scientific) fitted with the Acclaim PepMap RSLC column (Thermo Fisher Scientific), making use of a 60 minutes gradient (2–95% v/v acetonitrile, 0.1% v/v formic acid) running at a flow rate of 300 nl/minute. Peptides eluted from the nano LC column were subsequently ionized into the Q Exactive™ Plus mass spectrometer (Thermo Fisher Scientific). The electrospray source was fitted with an emitter tip 10μm (New Objective, Woburn, MA) and maintained at 1.5 kV electrospray voltage. The temperature of the capillary was set to 250°C. Precursor ions were selected for MS/MS fragmentation using a data-dependent “Top 10” method operating in FT-FT acquisition mode with HCD fragmentation. FT-MS analysis on the Q Exactive™ Plus was carried out at 70,000 resolution and an AGC target of 1×10^6^ ions in full MS. MS/MS scans were carried out at 17,500 resolution with an AGC target of 2×10^4^ ions. Maximum injection times were set to 30 and 50 milliseconds respectively. The ion selection threshold for triggering MS/MS fragmentation was set to 25,000 counts and an isolation width of 2.0 Da was used to perform HCD fragmentation with normalised collision energy of 27.

### Bioinformatics and statistics

The raw files were searched using Proteome Discoverer 2.4 software (Thermo Fisher Scientific) incorporating the Sequest search algorithm employing the *Homo sapiens* Uniprot FASTA databases. Peptide identifications were determined taking into account a 20-ppm precursor ion tolerance and 0.1 Da MS/MS fragment ion tolerance for FT-MS and HCD fragmentation respectively. Peptide modifications were also considered whereby cysteine carbamidomethylation was considered a static modification. Variable modifications included methionine oxidation, asparagine and glutamine deamidation, lysine biotinylation, and acetylated N-terminal residues. Trypsin was set as the enzyme of use, allowing for three missed cleavages at the most. Data was also processed using a label-free quantitation (LFQ) workflow employing the Minora Feature node, making use of a Protein FDR validator node which estimates the false discovery rates at the protein level as well as a percolator node to estimate the FDR at the PSM level. Results were adjusted so that the final global FDR was less than 1% at the protein and peptide level. A *q*-value of 0.01 was required to validate protein identifications.

Statistical analyses were typically conducted using GraphPad Prism 8.2.1 software or Ingenuity Pathway Analysis (IPA). In GraphPad Prism, statistical analyses involved the use of a paired t-test. Comparisons were considered significant if the *p*-values were less than 0.05.

Statistically significant protein functions were identified using Ingenuity Pathway Analysis (IPA). Here a Right-Tailed Fisher’s Exact Test was used. Results were considered statistically significant if the *p*-value was less than 0.05.

### Adeno-associated viruses (AAV)

Vectors encoding full length human WT cyclin F or cyclin F carrying the S621G mutation (n-terminal V5-tagged) was cloned into a rAAV vector under the human synapsin promoter using the plasmid pAAV-hSyn-EGFP (gift from Bryan Roth; Addgene, #50465) as backbone and removing EGFP. The same vector with EGFP expression was used as control. Packaging of rAAV9 vectors were performed as previously described (*37*) using the capsid AAV9.PHP.B (*38*). Briefly, HEK293T cells in 15cm dishes were each transfected with 12.5µg of vector plasmid containing gene of interest, 25µg of pFΔ6 and 12.5µg of AAV rep-cap using PEI-max in IMDM (Sigma-Aldrich). Cells were harvested 48 hours post-transfection by scraping and centrifuged at 350×g for 30 minutes. The cell supernatant was subjected to polyethylene glycol (PEG) precipitation and cell pellet was further lysed using a freeze-thaw cycle and combined with the PEG mixture. After lysis with sodium deoxycholate and 3 rounds of freeze/thaw cycles, the supernatant was collected for purification in an OptiSeal tube (Beckman-Coulter) containing iodixanol layers (15%, 25%, 40%, 54%; Sigma-Aldrich). Purified virus was collected using a 19G syringe, inserted just below the 405 gradient and during dialyzed and concentrated using Amicon Ultra-15 CFU with 100 kDa cutoff filter (Millipore). The virus was sterile filtered through a Spin-X 0.22 μm centrifuge filter (Corning).

### AAV transduction into primary neurons

For AAV transduction, cortical neurons were transduced with WT CCNF, CCNF S621G or EGFP AAV at MOI of 5000 on DIV 3. At 10 days *in vitro* (DIV), cells were treated with 0.2 µM of proteasome inhibitor, MG132, for 12 hours.

### Immunohistochemistry and microscopy

Post-mortem paraffin-embedded cervical spinal cord sections from ALS patients (n=4) and controls (n=3) were obtained from the New South Wales Brain Bank Network. For immunohistochemical staining, tissue sections were heated at 70°C for 30 minutes, deparaffinized with xylene and rehydrated with a descending series of ethanol washes. To retrieve antigens, sections were boiled for 20 minutes in low pH buffer (pH 6.1; Dako, CA, USA). Endogenous peroxidase activity and non-specific binding were blocked by incubation with 3% hydrogen peroxide in methanol for 15 minutes followed by 5% normal goat serum (Vector Laboratories, CA, USA) with 0.1% TWEEN-20 in PBS for 1 hour. Sections were incubated at 4°C overnight with primary antibody rabbit anti-RBM14 (1:100, Sigma-Aldrich) and then at room temperature for 1 hour with biotinylated goat anti-rabbit IgG (Vector Laboratories). The avidin-biotin complex detection system (Vector Laboratories) with 3,3’-diaminobenzide as chromogen (Dako) was used to detect the immunoreactive signal. Nuclei were counterstained with hematoxylin before sections were dehydrated with an increasing series of ethanol washes followed by xylene. Sections were coverslipped using Di-N-Butyle Phthalate in xylene (DPX, Dako).

Tissue sections were visualized using the ZEISS Axio Imager 2 microscope and analysed using Fiji Image J. To quantify RBM14 neuronal nuclei staining, each image was first deconvoluted with the Image J ‘H DAB’ Deconvolution Macro (*39*). A region of interest (ROI) was drawn around the neuron nucleus and pixel intensity was scored using the IHC Profiler plugin categorizing overall RBM14 staining in the ROI as either high positive, positive, low positive or negative (*40*). Only neurons with a clear nucleus were included. The number of neurons with RBM14 expression in the high positive and positive zone or low positive and negative zone were plotted and analysed with Fishers’ exact test (significance set to 0.0166667 after Bonferroni correction).

## RESULTS

### BioID identifies known and novel protein interaction partners of cyclin F

In order to identify transient and low abundance interaction partners of cyclin F, we used a proximity-based biotinylation method known as BioID (*31*). Here, we first cloned cyclin F in frame with a modified biotin ligase (denoted BirA*-cyclin F^WT^). In addition, we cloned a mutant of cyclin F^LP/AA^ (which has previously been reported to stabilize the interaction between cyclin F and transiently interacting proteins) in frame with BirA*, generating BirA*-Cyclin F^LP/AA^ (Figure 1a). In order to tightly control the expression of the fusion protein in cultured cells, we generated stably transfected HEK293 T-Rex Flp-In cell lines (Thermo). The T-Rex Flp-In system was selected as it ensures that only a single copy of the transgene is placed within the exact same insertion site in the host genome, whilst the expression of cyclin F, a cell cycle regulator, is controlled in a tetracycline-dependent manner.

**Figure 1.**
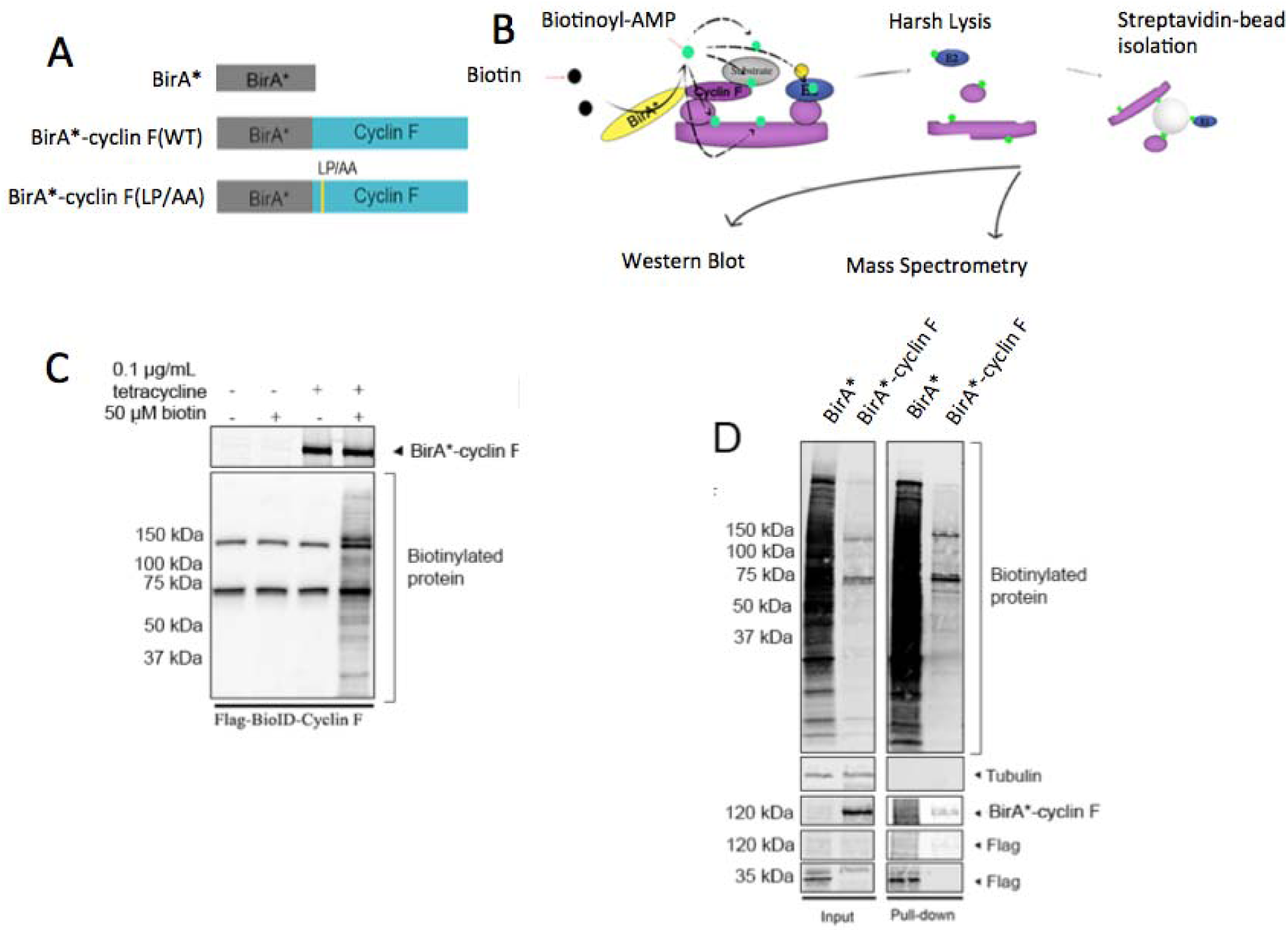
Identifying cyclin F interaction partners using BioID. **A.** BirA* alone, BirA*-cyclin F(WT) and BirA*-cyclin F(LP/AA) were stably transfected into Flp-In T-Rex HEK293 cells. **B.** Schematic showing process of biotinylation by BirA*, isolation of biotinylated proteins and identification by western blotting or mass spectrometry. **C.** Stably transfected HEK293 Flp-In cells were treated with tetracycline to induce gene expression. Addition of biotin led to the biotinylation of proteins in proximity to cyclin F, as detected by immunoblotting with fluorescently-tagged streptavidin (LiCor). **D.** The biotinylation profile of BirA* alone or cyclin F-BirA* before and after streptavidin-coated bead pull-downs.

To conduct the BioID experiments, transgene expression was firstly induced in HEK293 Flp-In T-Rex cells using tetracycline. This is followed by addition of biotin to cell culture media for 24 hours. The resulting cells were lysed in harsh lysis buffer, before the biotinylated proteins were isolated using streptavidin-coated beads. These resulting proteins are then analysed by immunoblotting and mass spectrometry (Figure 1B). To begin BioID experiments that identify binding partners of cyclin F, we first confirmed that there was no leakage in either transgene expression or biotin-labelling, whilst initiation of cyclin F-BirA* expression and the addition of biotin leads to the biotinylation of proteins (Figure 1C). To identify proteins in proximity to cyclin F, cells expressing either cyclin F-BirA* or BirA* alone were expressed in HEK293 Flp-In T-Rex cells. Here, we induced expression of the transgene with tetracycline, added biotin, and after 24 hours cells were harvested. Biotinylated proteins were enriched using Streptavidin Magnetic Beads (Thermo) and prepared for immunoblotting and subsequent proteomic analysis. Notably, immunoblotting revealed that the biotinylation profile was greater in BirA* only expressing cells (Figure 1D), which correlated with higher expression levels of BirA* compared to BirA*-cyclin F. Thus, prior to MS analysis we adjusted the input of isolated biotinylated proteins accordingly (Supplementary Figure 1).

We carried out an in-gel trypsin digestion of the biotinylated proteins followed by liquid chromatography-mass spectrometry (LC-MS/MS). In total, 918 proteins were identified, with 163 proteins found in cyclin F-BirA* and cyclin F-BirA* (LP/AA) combined, but not when biotin ligase was expressed alone (Figure 2A). The list of protein identifications was further filtered such that proteins were considered interaction partners of cyclin F if they: *i.* increased at least two-fold when comparing BirA*-cyclin F to BirA* and *ii.* were present in at least 2 out of 3 biological replicates of BirA*-cyclin F expressing cells. The final list of high-confidence interactors of cyclin F yielded 119 proteins (presented in Supplementary Table 1). Within this list, we identified cyclin F and RING-box protein 1 (Rbx1). Rbx1 and cyclin F are both essential units of the Skp1-Cul1-Fbx^(cyclin F)^ E3 ubiquitin ligase complex, confirming that the BioID assay has biotinylated proteins within close proximity as expected. Casein kinase II (CKII), another previously identified binding-partner of cyclin F (*26, 41*), was also found in this list.

**Figure 2.**
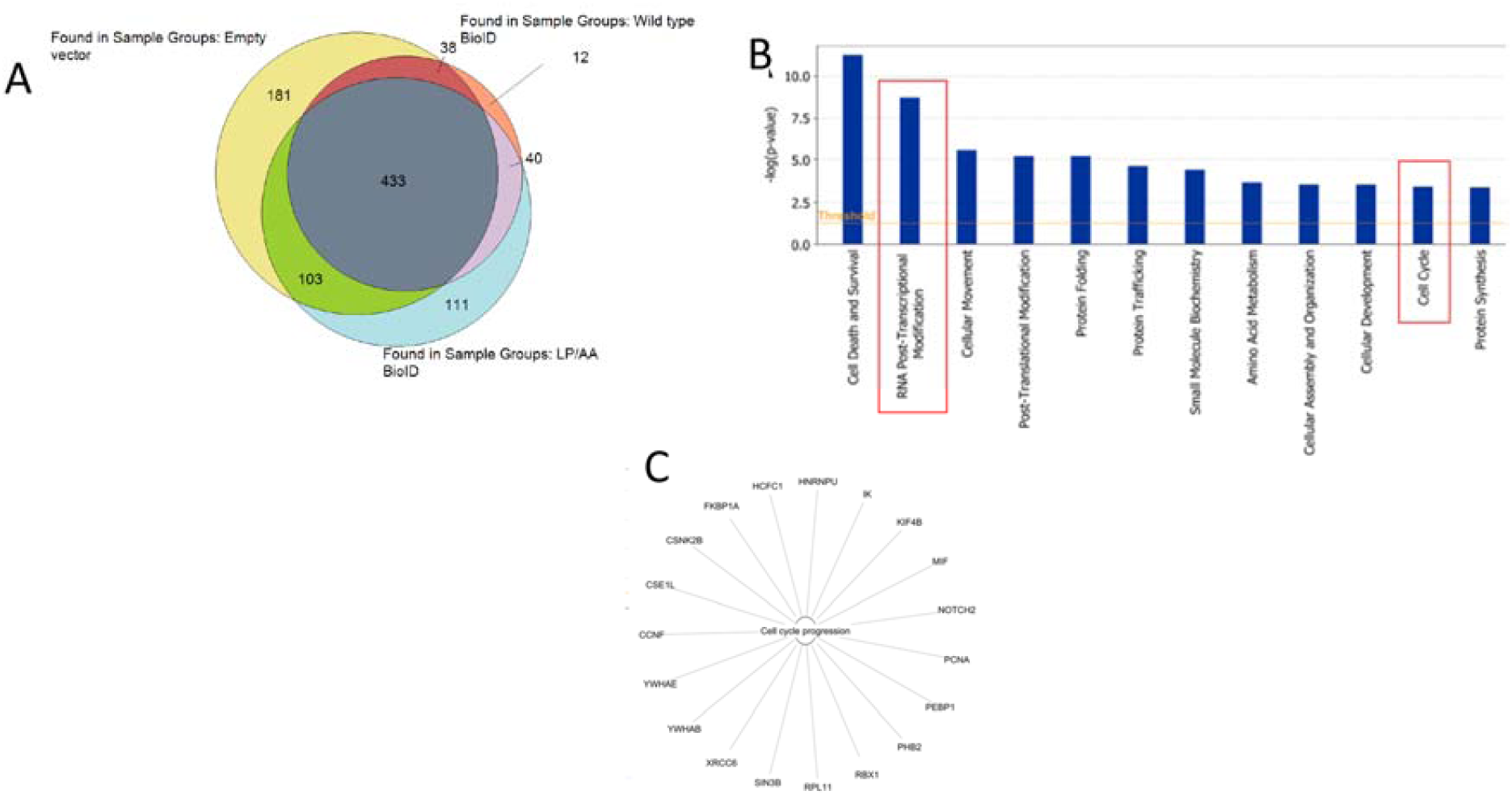
**A.** Proteomic analysis identified common and unique proteins biotinylated by BirA* alone, BirA*-cyclin F(WT) and BirA*-cyclin F(LP/AA). **B.** Ingenuity Pathway analysis (IPA) of protein interaction partners showing top twelve statistically enriched molecular processes for cyclin F interaction partners. **C.** Proteins interaction partners involved in cell cycle progression.

### Bioinformatic pathway analysis identifies known and novel functions of cyclin F

Next, Ingenuity Pathway Analysis (IPA) analysis was used to assign statistically significant molecular functions to proteins associated with cyclin F (Figure 2B). Consistent with known function of cyclin F, IPA unveiled a statistical enrichment of interaction partners with roles in ‘Cell Cycle Progression’ (p<3.51E-04) (Figure 2C), as well as ‘DNA Replication, Recombination and Repair’ (p<1.59E-02). Several novel molecular functions were also identified, one of which involves ‘RNA Post-Transcriptional Modification’ with specific functions of ‘Processing of mRNA’ (p=4.92E-07), ‘Splicing of mRNA’ (p=1.41E-06), ‘Processing of rRNA’ (p=1.82E-04), ‘Unwinding of mRNA’ (p=1.56E-02), ‘Annealing of hnRNA’ (p=1.56E-02), ‘Processing of RNA’ (p=1.90E-09) and ‘Splicing of RNA’ (p=2.70E-07).

We also used IPA to analyse the 111 proteins that were uniquely associated with cyclin F^LP/AA^, as this protein list includes stabilized interaction partners. In this list, we found a series of proteins which had roles in ‘RNA Damage and Repair’ (p=4.27E-10), ‘Processing of RNA’ (p=1.38E-08), ‘Processing of rRNA’ (p=7.98E-08), ‘Splicing of RNA’ (1.83E-04). Within the list of proteins with roles known to be involved in RNA metabolism, there were also a series of RNA binding proteins including TAF15, EWS and RBM14; proteins also known to form paraspeckles (Figure 3). Notably, IPA predicted changes in the homeostasis of these proteins to affect the expression of RNA, prompting further investigation into the relationship between cyclin F and these interaction partners.

**Figure 3.**
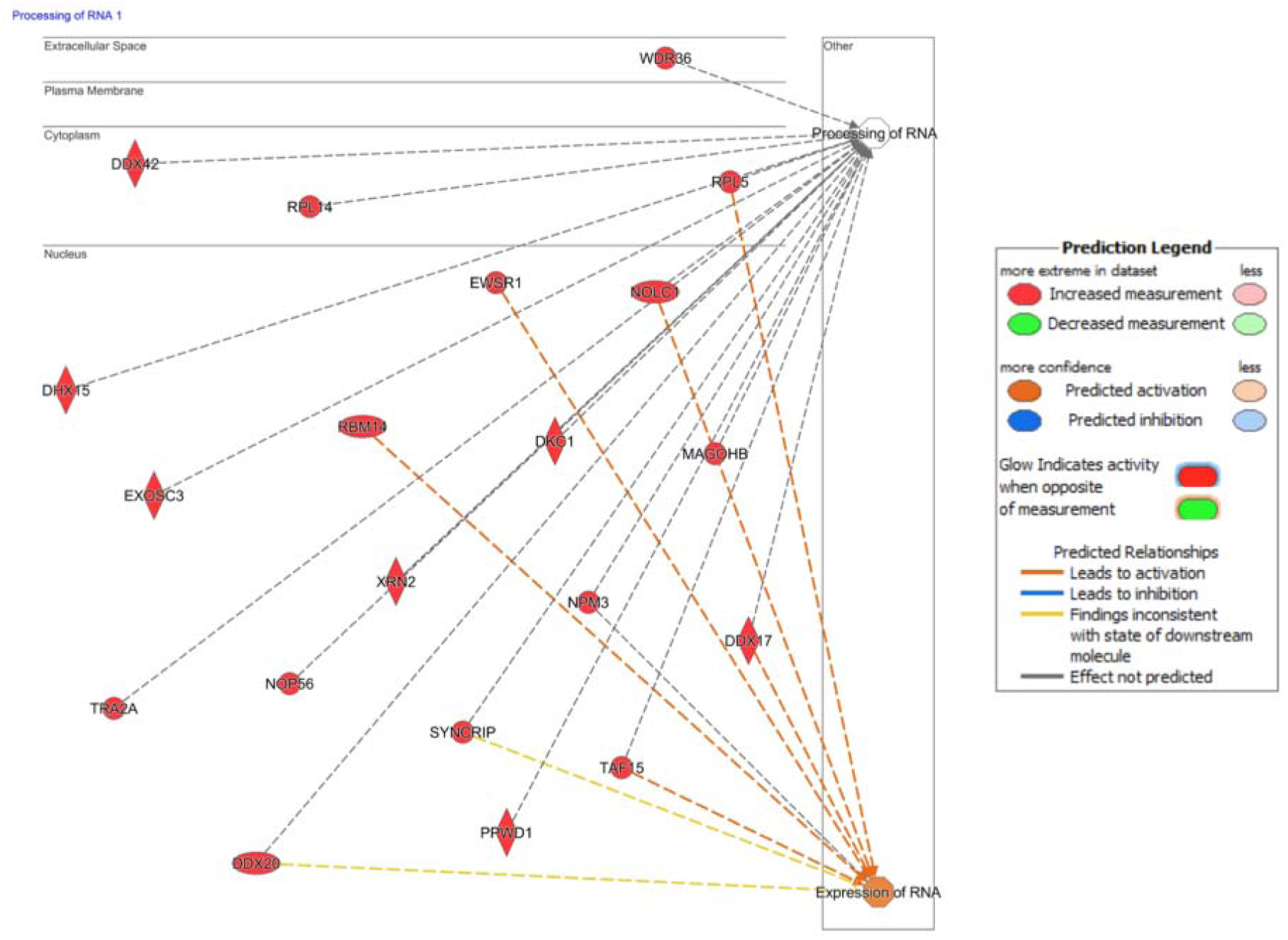
Protein interaction partners involved in RNA processing and expression along with subcellular localisation.

### Cyclin F is closely associated with paraspeckle proteins

Given that RBM14 is essential for building subnuclear paraspeckles (*42*), we validate the interaction between cyclin F and RBM14 using standard immunoprecipitation and immunoblotting (Figure 4A). Here we noted that cyclin F(WT)-flag and cyclin F(S621G)-flag could co-immunoprecipitate with endogenous RBM14, with no effect from the mutation. In addition, both overexpressed cyclin F(WT)-flag and cyclin F(S621G)-flag could co-immunoprecipitate with RBM14-HA. Conversely, RBM14-HA could co-immunoprecipitate with both cyclin F(WT)-flag and cyclin F(S621G)-flag (Figure 4B).

**Figure 4.**
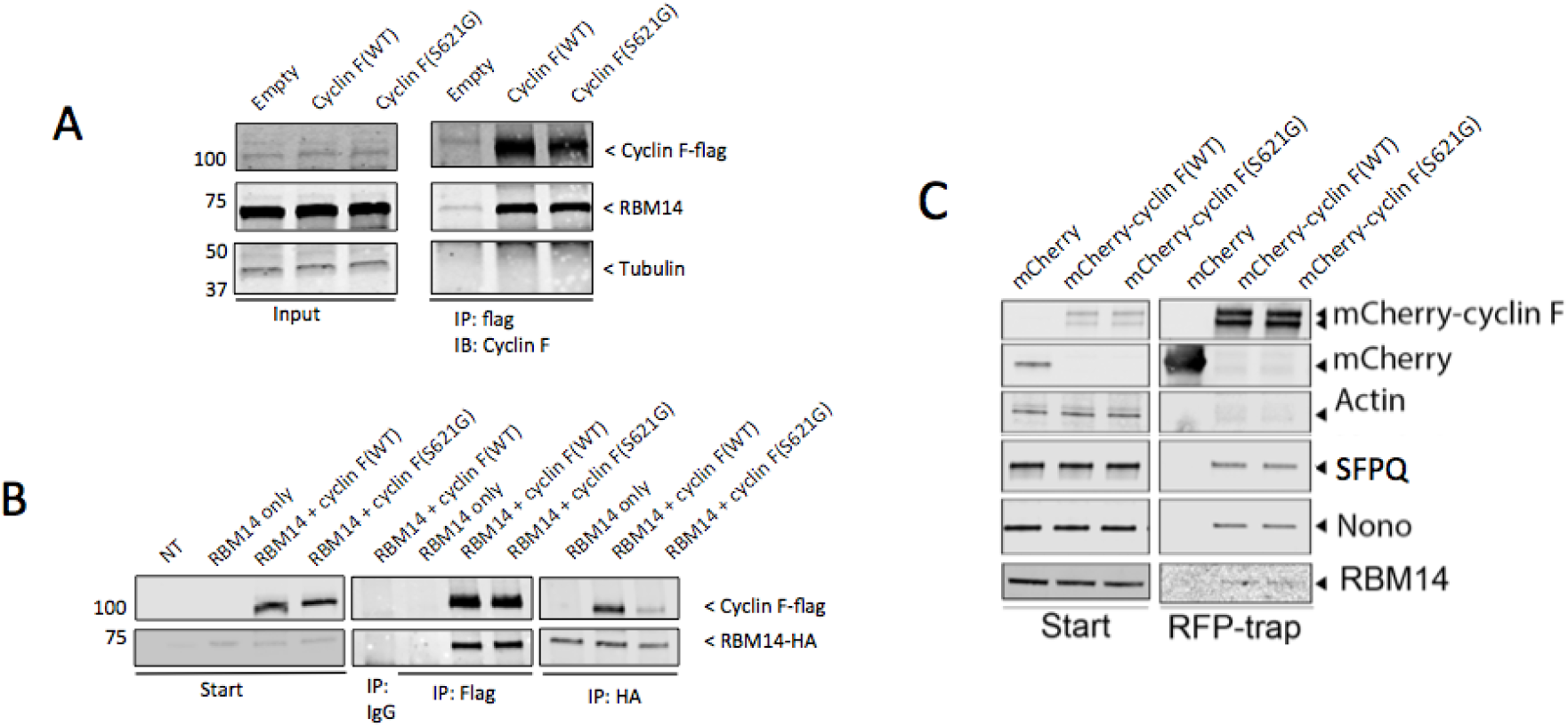
Cyclin F interacts with paraspeckle proteins. **A**. Flag-tagged cyclin F(WT), cyclin F(S621G) or an empty vector control were transfected into HEK293 cells. Anti-flag antibody was used to immunoprecipitate flag-tagged proteins. Eluates were evaluated by immunoblotting with the antibodies specified. **B**. Flag-tagged cyclin F(WT) or cyclin F(S621G) were co-transfected alongside RBM14-HA in HEK293 cells. Anti-flag or anti-HA antibody was used to immunoprecipitate flag-tagged or HA-tagged proteins as specified. Eluates were evaluated by immunoblotting with the antibodies specified. **C.** mCherry-cyclin F(WT) or mCherry-cyclin F(S621G) were transfected into HEK293 cells. An RFP-trap was incubated with lysates to immunoprecipitate mCherry-tagged proteins. Eluates were evaluated by immunoblotting using the antibodies specified.

Next we questioned whether cyclin F could also bind other essential components of the paraspeckle complex, NONO and SFPQ. Indeed, mCherry-cyclin F was found to immunoprecipitate with these essential paraspeckle components in addition to RBM14 (Figure 4C). In all cases, both cyclin F^WT^ and cyclin F^S621G^ were able to co-immunoprecipitate with these paraspeckle components. Together the data validated the mass spectrometry data and further demonstrated that cyclin F interacted with protein components of the paraspeckle complex.

### Cyclin F^S621G^ causes disruption of paraspeckle homeostasis in primary neurons

Previously it has been established that proteasome inhibitor, MG132 is able to initiate paraspeckle assembly and lead to elongation of the paraspeckle structure. During this time, paraspeckle protein levels remain largely consistent (*43*). To assess the effect of cyclin F^S621G^ on paraspeckle regulation in response to MG132, we overexpressed cyclin F^WT^, cyclin F^S621G^ or an empty vector in primary mouse cortical neurons by AAV infection, then collected protein lysates following treatment with 0.2 µM of MG132 or a vehicle control (Figure 5A). Notably we observed that, in response to MG132 treatment, there was a significant increase in RBM14 of 1.54 fold (Figure 5B) as well as a significant increase in SFPQ levels of 1.21 fold-change (Figure 5C) in cyclin F^S621G^ overexpressing cells compared to the wild-type control. There was no significant difference in the expression of NONO in response to MG132 treatment (Figure 5D) suggesting that mutant cyclin F may lead to a disruption in the homeostasis of some essential paraspeckle components.

**Figure 5.**
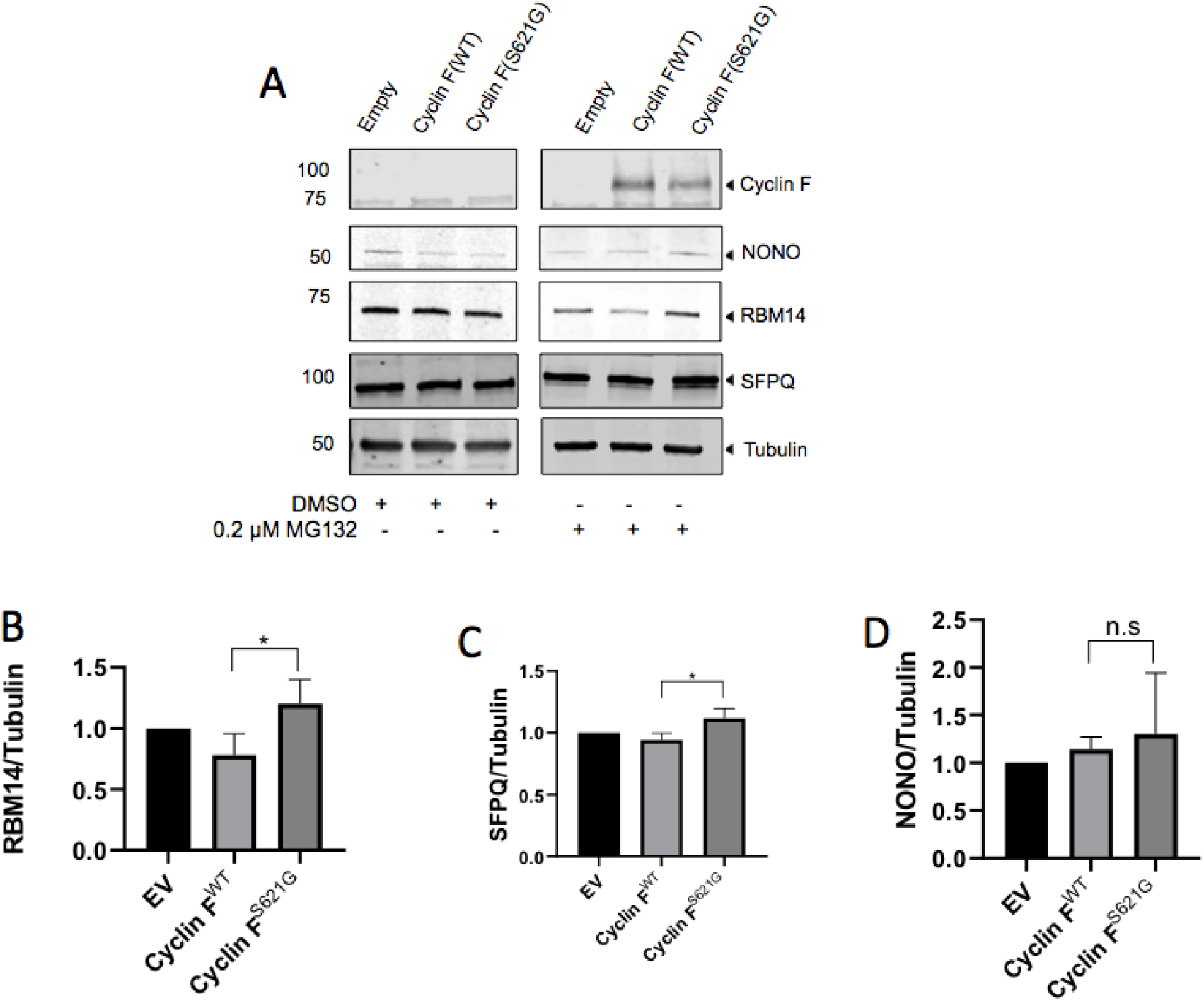
Cyclin F^S621G^ causes defective turnover of paraspeckle components in primary neurons. **A**. Primary neurons were transduced using constructs encoding cyclin F^WT^, cyclin F^S621G^ or an empty vector control. Transduced neurons were treated with 0.2 µM MG132 or a vehicle control for 24 hours before cells were lysed in RIPA buffer and analysed by immunoblotting using the antibodies as indicated. **B**. Densitometry of RBM14 upon MG132 treatment normalised to Tubulin (n=4, *:*p*<0.05). **C**. Densitometry of SFPQ upon MG132 treatment normalised to Tubulin (n=4, *:*p*<0.05). **D**. Densitometry of NONO upon MG132 treatment normalised to Tubulin (n=4, n.s; not significant).

### RBM14 homeostasis is dysregulated in the motor cortex and spinal cord of ALS patients

RBM14 homeostasis has not previously been reported in patient postmortem tissue. To determine whether RBM14 levels are dysregulated in post-mortem ALS patient tissue, we measured RBM14 levels in motor cortex and spinal cord neurons via semi-quantification of immunohistochemical labeling. Since RBM14 is a known nuclear protein, we specifically compared RBM14 expression in neuronal nuclei from control and ALS patients (Table 2). In control spinal cord neurons, RBM14 showed either primarily nuclear staining or staining in both nuclei and cytoplasm. In comparison, in ALS patient tissues, there was a significant reduction of nuclear RBM14 in spinal cord neurons (Figure 6).

**Figure 6.**
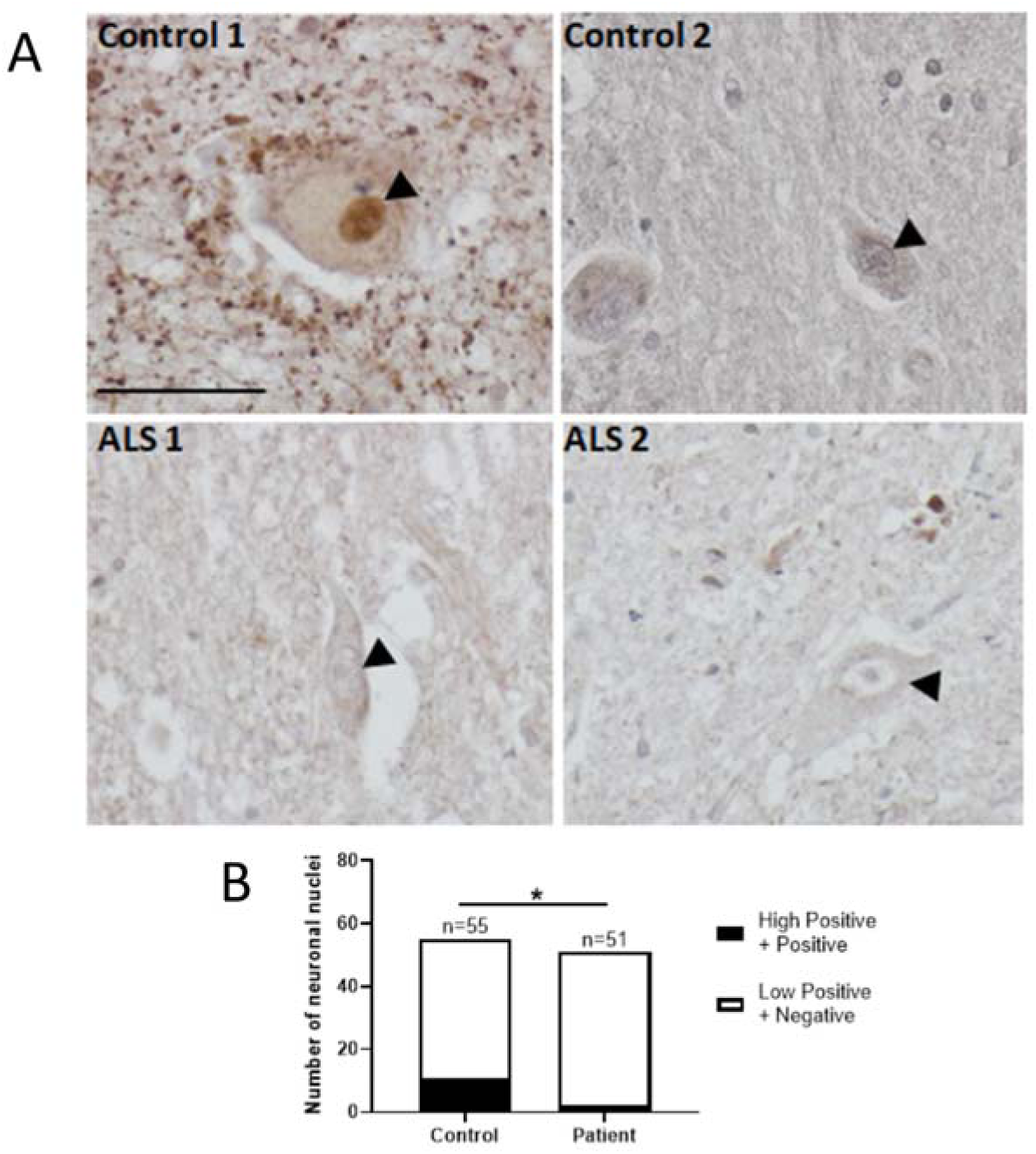
RBM14 is reduced in the brain and spinal cord of ALS patients. **A.** Representative images of RBM14 immunohistochemical staining in control (n=3) and ALS patient (n=4) spinal cord tissues. Arrowheads indicate RBM14 staining in neuronal nuclei. Scale bar=50 µm. **B.** Semi-quantification of RBM14 IHC staining showed a significant reduction of RBM14 in neuronal nuclei from ALS patients compared to controls. A total number of 55 control neurons and 51 patient neurons were analysed using Image J IHC Profiler Plugin (Fisher’s exact test, *: *p*<0.05).

## DISCUSSION

In this study, we have identified novel protein interactors of cyclin F using BioID coupled with mass spectrometry. We found that cyclin F was closely associated with paraspeckle proteins including RBM14, NONO, and SFPQ. Furthermore, we demonstrate that the homeostasis of RBM14 and SFPQ, essential components of the paraspeckle complex, is influenced by cyclin F and becomes defective when cyclin F carries an S621G mutation linked to ALS/FTD. Finally, we show yet another paraspeckle protein, RBM14, may be involved in ALS pathogenesis through the dysregulation of protein levels in post-mortem motor cortex and spinal cord of ALS patients.

We previously reported the identification of disease-causing variants in *CCNF* in familial and sporadic ALS/FTD patients and have reported defects in major protein degradation systems in cells overexpressing cyclin F(S621G) (*1*). Given these known deficits, it is logical to predict that the role of cyclin F in ALS pathogenesis may be associated with defective protein degradation pathways. Given that many interaction partners of cyclin F were unknown, we undertook unbiased proteomic screening to identify interaction partners cyclin F. To take advantage of the T-Rex and Flp-In systems (allowing controlled copy number integration at the exact same site) we performed the BioID assay in HEK293 cells, and in doing so, identified more than 100 high-confidence interaction partners. We acknowledge that many of these may not be relevant to motor neurons as their biological function is related to activities such as cell division. However, we did identify several proteins involved in RNA processing pathways that are likely to be relevant to ALS/FTD.

The BioID assay identified a close association of cyclin F with a group of paraspeckle proteins. Further work revealed that an ALS-causing mutation in cyclin F leads to the defective homeostasis of essential paraspeckle proteins, RBM14 and SFPQ. Paraspeckles are a class of subnuclear bodies that form within the interchromatin space of mammalian cells (*44*). These RNA-protein structures form as RNA binding proteins interact with the long non-coding RNA (lncRNA), *NEAT1* (*45*). Alterations to paraspeckle assembly and function has important implications in the context of neurodegeneration as paraspeckles have a clear role in controlling gene expression (*44*). In particular, paraspeckles are known to regulate multiple cellular processes such as cell stress responses, cellular differentiation and viral infections (*44*). Therefore, disruption to paraspeckle assembly or function results in inability to rapidly transcribe stress-responsive proteins required for maintaining cellular viability. Notably, the formation of paraspeckles, and the dysregulation of this process, is emerging as a biological marker of ALS. For example, the assembly of paraspeckle proteins around *NEAT1_2* has been reported in spinal motor neurons of early-stage ALS patients (*46*). In addition, compromised paraspeckle formation has been identified in cell and animal models of FUSopathies, with mislocalised FUS resulting in neuronal inclusions of paraspeckle components (*47*). In both studies, the increased levels of paraspeckle assembly may represent a downstream, protective cellular response to stress. We now report a different type of possible involvement of paraspeckles in ALS pathogenesis. We show that RBM14 homeostasis is dysregulated in post-mortem brain and spinal cord of ALS patients. RBM14 has been shown to connect key paraspeckle subcomplexes, a function which requires the presence of its prion-like domain (*42*). Thus, the dysregulation of RBM14 (and potentially other paraspeckle proteins), a core paraspeckle protein, may impair paraspeckle assembly/function and leave motor neurons vulnerable to cellular stress and therefore more susceptible to neurodegeneration. Importantly, in this study, we have shown that RBM14 dysregulation occurs in the brain and spinal cord of patients, regardless of the presence of *CCNF* mutation, suggesting that the dysregulation of RBM14 homeostasis may be one contributing step in the multi-stage pathogenesis of ALS.

Of the more than 30 genes (and their corresponding protein products) that are now linked to ALS, two broad functional categories have emerged; protein-degradation pathways (indirectly because of the presence of abnormal protein aggregates, and directly through regulators of protein degradation such as cyclin F and ubiquilin-2) and RNA processing. However, the link between these two distinct groups of proteins remain poorly understood. Perhaps the strongest association to date is represented by TDP-43 and to a lesser extent FUS, which are both major constituents of intraneuronal aggregates, and their core function being associated with RNA processing (*48*). We now present a new and different linkage, demonstrating that cyclin F influences the homeostasis of key paraspeckle components. Future studies should look to identify the cellular processes, such as RNA processing in response to stress stimuli, that may become dysregulated due to the reduction of RBM14 in the nucleus of affected motor neurons, as this may provide deeper insight into the underlying causes of ALS/FTD. Notably, this study also identifies RBM14 dysregulation in sporadic cases of ALS. Together the data further link the dysregulation of the ubiquitin-proteasome system and RNA processing to the pathogenesis of ALS/FTD.

## CONCLUSIONS

This study employed an unbiased proteomic screening assay which revealed that cyclin F interacts with several core proteins of the paraspeckle complex. Using immunoprecipitation, we confirmed the interaction between cyclin F and three paraspeckle proteins; RBM14, NONO and SFPQ. Notably, we demonstrate that the pathogenic cyclin F^S621G^ variant disrupts the homeostasis of these proteins and their responsiveness to a stressor that stimulates paraspeckle formation. Finally, we report for the first time that RBM14 levels are dysregulated in brain and spinal cord of ALS patients relative to healthy patient controls. Collectively, these data suggest that cyclin F may influence stress responses through modulation of the paraspeckle complex, and that disruption in paraspeckle homeostasis may contribute to the molecular pathogenesis of ALS/FTD.

**Table 1.** Details of patient tissue

| **Case** | **Tissue type** | **Age**<br>**(y)** | **Gender** | **PMI (hr)** | **Disease**<br>**onset** |
| --- | --- | --- | --- | --- | --- |
| *Control* | Motor cortex | 37 | Male | 24 | N/A |
| *Control* | Spinal cord and motor<br>cortex | 61 | Male | 30 | N/A |
| *Control* | Spinal cord and motor<br>cortex | 80 | Male | 12 | N/A |
| *Control* | Spinal cord | 75 | Male | 34 | N/A |
| *SALS* | Motor cortex | 81 | Male | 70 | 80 |
| *SALS* | Motor cortex | 84 | Male | 23 | 74 |
| *FALS (C9orf72)* | Motor cortex | 60 | Male | 99 | 58 |
| *FALS (Unknown*<br>*mutation)* | Spinal cord and motor<br>cortex | 54 | Female | 16 | 53 |
| *SALS (C9orf72)* | Spinal cord | 65 | Female | 18 | 65 |
| *SALS* | Spinal cord | 71 | Male | 12.5 | 70 |
| *FALS (C9orf72)* | Spinal cord | 75 | Male | 74 | 21.5 |

## Abbreviations

AAV: Adeno-associated viruses
ACN: Acetonitrile
AGC: Automatic gain control
ALS: Amyotrophic Lateral Sclerosis
AmBic: Ammonium bicarbonate
BCA: Bicinchoninic acid assay
BioID: Proximity dependent *Bio*tin *Id*entification
Bis-Tris: 1,3-bis(tris(hydroxymethyl)methylamino)propane
BSA: Bovine serum albumin
DIV: Days *in vitro*
DMEM: Dulbecco’s Modified Eagle Medium
DMSO: Dimethyl sulfoxide
DNA: Deoxyribonucleic acid
DTT: Dithiothreitol
EDTA: Ethylenediaminetetraacetic acid
EGFP: Enhanced green fluorescent protein
FA: Formic acid
FT-MS: Fourier transform-mass spectrometry
FBS: Fetal bovine serum
FDR: False discovery rate
FTD: Frontotemporal Dementia
HCD: Higher energy collision dissociation
MS: Mass spectrometry
IAA: Iodoacetamide
IgG: Immunoglobulin G
IMDM: Iscove’s Modified Dulbecco’s Medium
IP: Immunoprecipitation
IPA: Ingenuity Pathway Analysis
LFQ: Label-free quantitation
MND: Motor neurone disease
MOI: Multiplicity of Infection
nanoESI: Nanoelectrospray ionization
NHEJ: Non-Homologous End Joining
NP-40: Nonidet P-40
PBS: Phosphate buffered saline
PBST: Phosphate buffered saline containing Tween-20
PEG: Polyethylene glycol
PEI: Polyethylenimine
PSM: Peptide-spectrum match
RIPA: Radioimmunoprecipitation assay buffer
ROI: Region of interest
SCF: Skp1-Cul1-F-Box
SDS: Sodium dodecyl sulphate
SDS-PAGE: Sodium dodecyl sulphate polyacrylamide gel electrophoresis
UPS: Ubiquitin-proteasome system

## DECLARATIONS

### Ethics approval and consent to participate

International, national, and/or institutional guidelines for the care and use of animals were followed. Ethics approval was also obtained for the use of human tissue.

### Consent for publication

Not applicable

### Availability of data and material

The mass spectrometry proteomics data have been deposited to the ProteomeXchange Consortium via the PRIDE partner repository with the dataset identifier PXD014163 and 10.6019/PXD014163.

### Authors’ Contributions

R.C. and A.L. and S.L.R. conceptualized the project. S.L.R. conducted the BioID studies, MS analysis, follow-up biochemical studies and wrote the manuscript. F.C. assisted with MS sample runs. A.D. assisted with stable cell line generation. S.Y. and N.G. generated lysates from patient tissue and conducted IHC studies using patient tissue. C.G.A. and Y.D.K. conducted primary neuron transduction and drug treatment. M.P.M., I.B., A.L., R.C., J.M.D., M.M., L.M.I., B.S. assisted with editing the manuscript. All authors read and approved the final manuscript.

## Acknowledgements

This research was supported by access to the Australian Proteomics Analysis Facility (APAF) established under the Australian Government’s NCRIS program.

