## Supplementary Figure 1 for "BioID analysis of the cyclin F interactome reveals that ALS-variant cyclin F alters the homeostasis of paraspeckle-associated proteins"

Supplemental Figure 1: Coomassie stained gel showing input of biotinylated proteins used for Mass Spectrometry analysis.

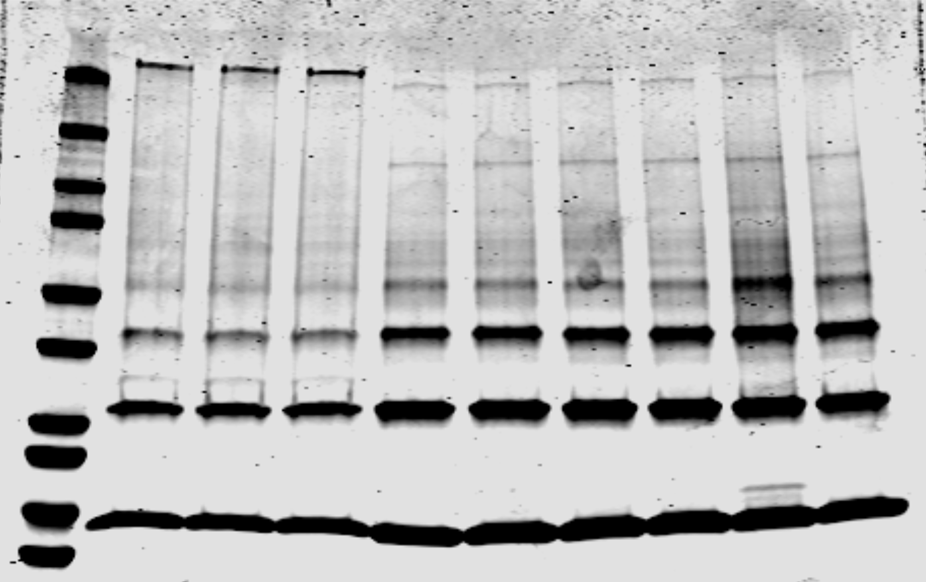

BirA*

BirA*

BirA*

BirA*-cyclin F

BirA*-cyclin F

BirA*-cyclin F(LP/AA)

BirA*-cyclin F(LP/AA)

BirA*-cyclin F(LP/AA)

BirA*-cyclin F

75

100

150

250

50

37

25

20

15

10

kDa
